# Functional definition of thyroid hormone response elements based on a synthetic STARR-seq screen

**DOI:** 10.1101/2022.02.23.481571

**Authors:** Frédéric Flamant, Romain Guyot

## Abstract

When bound to thyroid hormone, the nuclear receptor TRα1 activates the transcription of a number of genes in many cell types. It mainly acts by binding DNA as an heterodimer with RXRs at specific response elements related to the DR4 consensus sequence. However, the number of DR4-like elements in the genome exceed by far the number of occupied sites, indicating that minor variations in nucleotides composition deeply influence the DNA-binding capacity and transactivation activity of TRα1. An improved protocol of synthetic STARR-seq was used to quantitatively assess the transcriptional activity of thousands of synthetic sites in parallel. This functional screen highlights a strong correlation between the affinity of the heterodimers for DNA and their capacity to mediate the thyroid hormone response.

## Introduction

Nuclear receptors mediate the action of a variety of natural and artificial ligands by regulating gene transcription (1). They bind DNA response elements by their N-terminal domain as homodimers or heterodimers. The binding of ligand in their C-terminal domain induces a conformational change, and, depending on the agonist or antagonist nature of the ligand, recruit transcription coactivators or corepressors. Response elements are composed of two half sites, which DNA sequence and arrangement dictate the type of dimers that can bind. Recent genome wide analyses revealed that, while there are more than 50 000 putative response elements in the genome for each nuclear receptor, only few thousands are actually occupied in chromatin. The number of genes which transcription is responsive to the cognate ligand is further limited to few hundreds. One hypothesis to explain this restricted response is that the DNA/nuclear receptor/ligand complex has allosteric properties (2,3). Under this hypothesis, the exact nucleotide sequence of the DNA response element indirectly would not only determine its affinity for receptor binding but also influence the capacity of the distant ligand binding domain to recruit transcription corepressors or coactivators. It is therefore important to provide a functional characterization of the response elements, in addition to assess their capacity to bind the nuclear receptors *in vitro*.

As an example, we selected the well-studied TRα1/RXRα heterodimer, which activates transcription when TRα1 bind the thyroid hormone (3,3′,5-triiodo-L-thyronine or T3) (4). *In vitro* studies (5) and motif analysis of ChipSeq datasets (6–9) previously established that this heterodimer preferentially binds the so-called DR4 element, in which two half-site are separated by a 4-nucleotides spacer (5’<u>AGGTCA</u>NNNN<u>AGGTCA</u>3’). Structural analysis indicates that the heterodimer orients so that RXR binds to the upstream half-site, and TRα1 to the downstream site. It also shows that the contact between DNA and TRα1 amino-acids extend to the last two nucleotides of the spacer (10). In order to assess the functional consequences of variations around this consensus, we performed a functional screen of variant DR4 elements, using an adaptation of the recently developed STARR-seq method (11,12).

## Material and method

### Library construction

A 105 nucleotides oligomer containing a degenerated DR4-element 5’ATACTAGTCGCACTACGATCCTGCCGGGTGG<u>NGGTCA</u>NNNN<u>RGGNNA</u>ATCCCCTCCCACACCTAATGCAGAGCTAGCCA3’ (DR4 half-sites are underlined) was amplified with the following primers: 5’GGCTAACCGGTGCTAGCATACTAGTCGCACTACGATC3’ and 5’TGAAAGTCGACGCTAGCTCTGCATTAGGTGT3’. The nucleotides flanking the degenerated consensus in the template oligonucleotide were copied from the flanking sequence of a DR4 element located 1520 nucleotides upstream to the transcription start site of *Hairless*, which is a well characterized TRα1 target gene (chr14:70552167-70552187 on the mouse genome; GRCm38/mm10 assembly). The resulting amplicon was purified using the Qiaquick purification kit (Qiagen), cut with the AgeI and SalI restriction enzymes and purified again. The cloning vector was prepared by cutting the hSTARR-seq-ORI plasmid (13) with the AgeI and SalI restriction enzymes. Extremities were dephosphorylated with rAPid Alkaline Phosphatase (Sigma-Aldrich) and the 2.5kb fragment was purified after agarose gel electrophoresis using the QIAquick Gel Extraction Kit (Qiagen). Sixty ng of this vector were ligated (T4 DNA ligase Biolabs) to 60 ng of the amplicon insert in 30 μl of buffer. The ligation product was used to transform XL10-Gold ultra-competent cells (Agilent). Library DNA was purified from the entire bacteria library grown on agarose plates. DNA was extracted from the scrapped colonies using the QIAprep Spin Miniprep Kit (Qiagen) and further purified after a phenol/CHCl3 extraction by ethanol precipitation.

### Plasmids construct

Reporter construct with luciferase reporter were based on the pGL4.12 [Luc2CP] (GB Acc# AY738224) (Promega) in which luciferase expression is driven by a minimum promoter derived from the HSV thymidine kinase gene. A single DR4 element was cloned upstream to this minimal promoter between the XhoI and BglII restriction enzymes.

### Cell transfection

HEK293 cells were seeded in 24 well-plates (10^5^ cells/well). Cells were transfected the following day with plasmid or library DNA using the TransIT-Lt1 transfection reagent (Mirus Corporation Madison WI, USA). T3 was added 6h later and cells were harvested the following day for either luciferase activity measurement or RNA analysis. Luciferase activity was measured with the luciferase assay reagent (Promega) and quantified with the Centro luminometer (Berthold Technologies). RNA was extracted using the Nucleospin RNA kit (Macherey Nagel). Contamining DNA was eliminated by two steps of DNase I treatment performed on the extraction column according manufacturer instructions. Before reverse transcription, an additionnal DNase I treatment was performed on 1μg of RNA using the RNase-Free DNase set (Qiagen). After 15 min incubation at room temperature, 25 mM EDTA was added and enzyme inactivated by 10 min incubation at 65°.

### Deep sequencing

RNA was reverse transcribed with Moloney Murine Leukemia Virus reverse transcriptase (Promega) and the DNA sequences corresponding to the 3’ end of the vector-driven transcript were amplified by PCR with 3 pairs of primers (Pair 1: 5’TCGTCGGCAGCGTCAGATGTGTATAAGAGACAGTCACTGGAGTTGTCCCAATTCTTG3’ + 5’GTCTCGTGGGCTCGGAGATGTGTATAAGAGACAGCGTCGACGCTAGCTCTGCAT3’; Pair 2: 5’TCGTCGGCAGCGTCAGATGTGTATAAGA GACAGNTCACTGGAGTTGTCCCAATTCTTG3’ + 5’GTCTCGTGGGCTCGGAGATGTGTATAAGAGACAGNCGTCGACGCTAGCTCTGCAT3’; Pair 3: 5’TCGTCGGCAGCGTCAGATGTGTATAAGAGACAGNNTCACTGGAGTTGTCCCAATTCTTG3’,+:5’GTCTCGTGGGCTCGGAGATGTGTATAAGAGACAGNNCGTCGACGCTAGCTCTGCAT3’) suitable to directly prepare an Illumina sequencing library (Illumina Nextera V2 index kit). A control was performed were reverse transcriptase was omitted to ascertain the absence of contaminating DNA. Amplicons were sequenced both by Sanger sequencing or by deep sequencing (Illumina Microkit V2 300 cycles Miseq). Paired-end reads contained in the .fastq files were merged (fastq-join - Galaxy Version 1.1.2-806.1) and converted to tabular format using FASTQ Groomer (Galaxy Version 1.0.4) and FASTQ to Tabular converter (Galaxy Version 1.1.5). Tab delimited table files were curated from sequences which differ from the expected sequences at the conserved positions around the DR4 elements (20/20 identity). A R-script was used to produce counts tables from the sequence tables.

### Statistics

The χ^2^-square test was used to detect significant changes in the observed distribution of nucleotides at specific positions, with Bonferroni correction. Specific response elements with differential response to T3 were identified using count tables and Deseq2 Galaxy Version 2.11.40.7 (14).

## Results

The STARR-seq method consists in inserting a putative enhancer in the 3’ non-coding sequence of an expression vector carrying a minimal transcription promoter. The abundance of the mRNA carrying the putative enhancer at its 3’ end then becomes an estimate of the enhancer activity. The method is sensitive, quantitative and can be adapted to screen libraries of putative hormone responsive enhancers and response elements (15). We used here the most recent version of the method, which overcomes several technical pitfalls (13). A synthetic DR4 consensus element with degenerated nucleotides was used to generate a plasmid library, which putatively contains 32768 different DR4 sequences (Figure 1a). We amplified the library and transfected HEK293 cells in duplicate with either 0.6μg, 1.2μg or 2.4μg of DNA. T3 (10^−8^M) was added to the culture medium of half of the transfected cells and RNA was extracted 36 hours later. We prepared cDNA and amplified the vector encoded RNA and perform a direct Sanger sequencing of the amplicons (Figure 1b), Tide DNA (16) was used to quantify signal intensity on the Sanger electrophoretograms and provides a first indication that T3 has an influence on the representation of the different nucleotides at degenerate positions.

**Figure 1:**
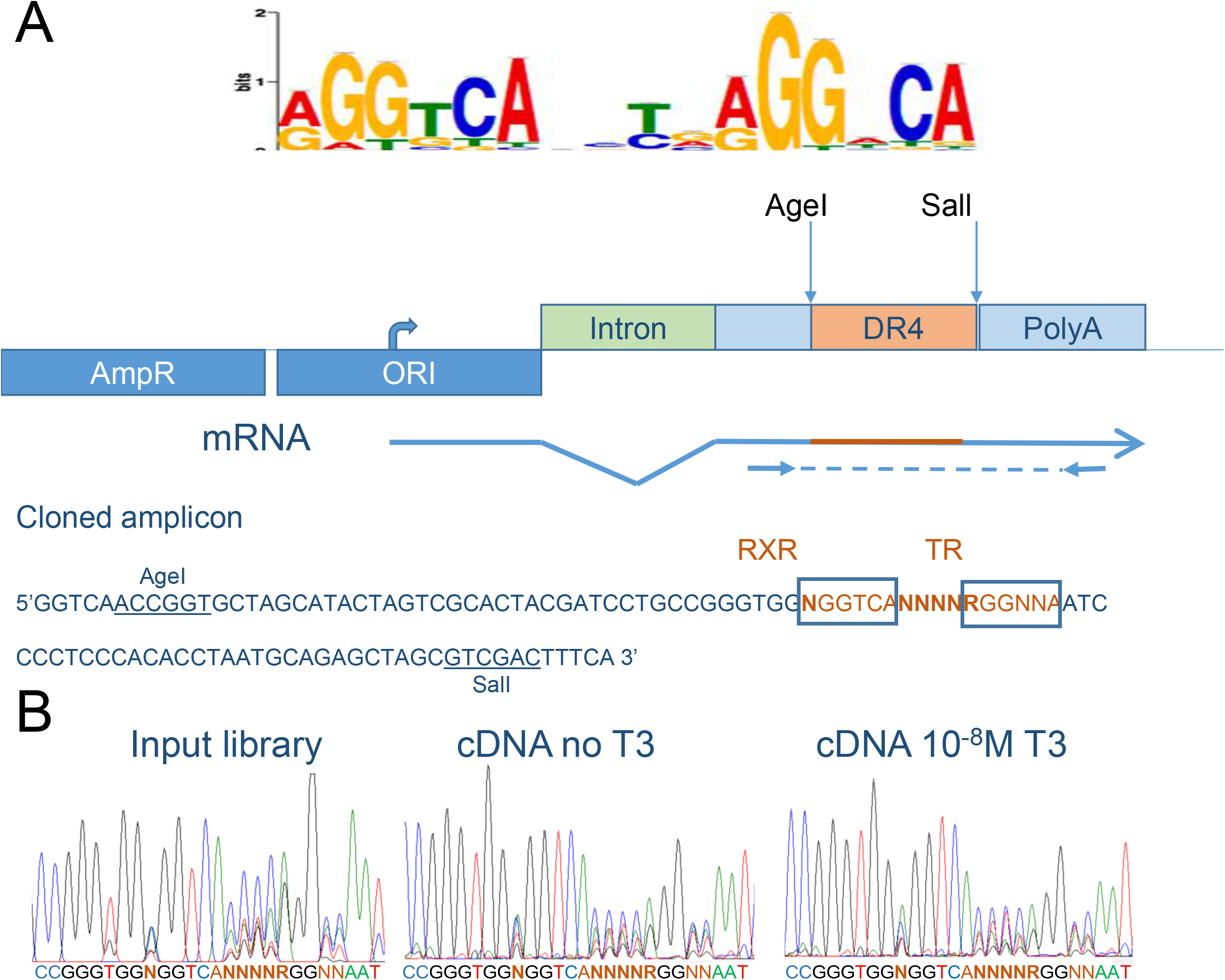
Library description. A) Logo of consensus DR4-like sequence discovered by *de novo* motif search (20) from a TRα1 ChIP-seq dataset obtained from mouse striatum neurons (9). According to structural data (10) the 5’half-site (5’RGGTCA3’) is occupied by RXR. The two 3’ nucleotides of the 4 nucleotides spacer and the 3’half-site (5’NNRGGNCA3’) contact TRα1. B. The expression vector is transcribed in eukaryotic cells from a cryptic minimal promoter present in the Ori sequence, spliced and contains the cloned amplicon. Sequences flanking the DR4 element with degenerated nucleotides are flanked by sequences present in the *Hairless* gene promoter. The small arrows indicate the position of the primer used for cDNA amplification. C. Sanger sequencing of the input library (left) indicates the presence of a mixture of different nucleotides at the expected positions. Sequencing of cDNA prepared from transfected cells (right) confirm the heterogeneity of the library and suggests that T3 has an influence on the respective abundance of different mRNA / DR4-like elements in the amplified cDNA.

We then performed deep sequencing for amplicons prepared from either the library DNA, or the 6 different cDNA. After curation for sequencing errors, the different DR4 sequences were counted. A total of 9129 different DR4 sequences were identified among 1078919 reads obtained for the plasmid library, with the expected balance between nucleotides at the variable positions (Table S1). Based on these 9129 sequences, which represent only 28% of the 32768 possible combinations, we calculated the frequency of each nucleotide at each variable position and addressed whether T3 can influence this distribution. A significant enrichment, indicating an influence on transactivation, was observed for specific nucleotides at each position, which was highly significant according to χ^2^-square tests. Since the enrichment was more pronounced when 1.2μg of DNA was transfected (Table S2), we concentrated the analysis on this experiment using the other datasets only to confirm the main results. The conclusion from this analysis is that there is a clear preference for specific nucleotides at several positions. In particular A or G at position 1, T or C at position 9, A at position 11, C at position 15 are clearly favorable to transactivation (Table 1).

**Table 1:** Influence of T3 on the frequency of nucleotide at variable positions. Relative frequency of a given nucleotide at the indicated position (with T3/without T3). Values above 1.1 are in bold characters.

|  |  |  |  |  |  |  |  |  |  |  |  |
| --- | --- | --- | --- | --- | --- | --- | --- | --- | --- | --- | --- |
| Position | 1 |  | 7 | 8 | 9 | 10 |  |  | 14 | 15 |  |
| Seq 5' to 3' | N | GGTCA | N | N | N | N | A/G | GG | N | N | A |
| A | **1.47** |  | 1.04 | 1.01 | 0.67 | 1.07 | 1.05 |  | 1.00 | 0.78 |  |
| C | 0.81 |  | 0.97 | 1.00 | **1.12** | 1.01 |  |  | 0.89 | **1.26** |  |
| G | **1.25** |  | 0.88 | 0.97 | 0.79 | **1.12** | 0.90 |  | 0.85 | 0.89 |  |
| T | 0.68 |  | **1.14** | 1.01 | **1.30** | 0.83 |  |  | **1.24** | 0.91 |  |

We then asked if nucleotides at each position operate independently or rather if distinct combinations of nucleotides are more favorable to promote T3 response. At the 3’end of the motif, the non-random distribution of dinucleotides enrichment was obvious (Figure 2A). We used χ^2^ test to ask whether the frequency of some dinucleotides deviates from the expected frequency, assuming that nucleotide changes act independently. The deviations did not originate from the oligonucleotide synthesis or library preparation as the same χ^2^ test did not reveal any significant bias in the input sequencing library. Although some dinucleotides combinations were clearly favored (Table S3), the most active dinucleotides were the ones predicted from the previous single nucleotide analysis (5’AC, 5’TC).

**Figure 2:**
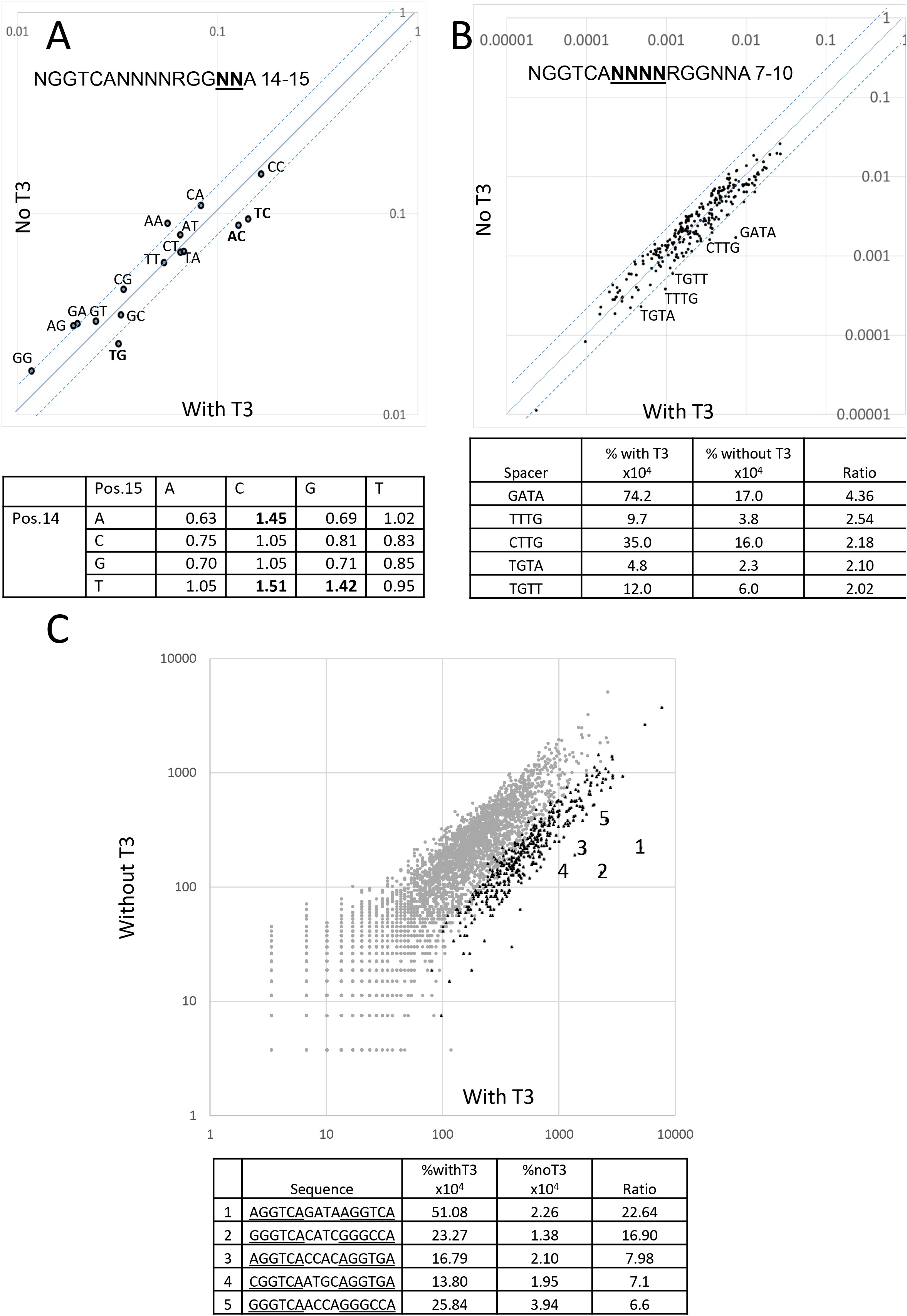
Features of the DR4 response element favoring T3 response. A. Frequency of specific dinucleotides in the downstream half-site contacted by TRα1 (positon 14 and 15). The table indicates the enrichment in cDNA library prepared from T3 treated cells compared to the input library. B Frequency of specific tetranucleotides in the spacer (positions 7 to 10). The table indicates the enrichment in cDNA library prepared from T3 treated cells compared to the input library for the 5 most favorable spacer tetranucleotides. C. Global analysis of all tested DR4 (Deseq2). Each grey dot represents a specific sequence. Black triangles are for the 455 sequences which are significantly enriched in the cDNA library prepared from T3 treated cells compared to the input library (adjusted p-value <0.05). The 5 most efficient DR4 sequences are indicated and listed in the table.

A similar analysis was applied to the spacer regions (Figure 2B) in which five combinations stand out as most favorable (fold-change >2), the 5’GATA sequence being the most enriched after T3 treatment. This result could not be predicted from the analysis of each position, as the expected frequency of this sequence, based on the individual nucleotides frequency, ranks at the 124 positions out of 256. χ^2^-square tests also showed that some combinations were overrepresented, with up to a 3-fold enrichment (Table S4).

We then used a more global analysis to address the influence of combining favorable nucleotides at variable positions to optimize the transactivation capacity of the DR4 element. Differential analysis using Deseq2 analysis (Figure 2C) showed that 455 of the tested sequences had a positive effect on mRNA level upon T3 treatment (Table S5). Among the sequences that were represented more than 100 times in each library, 5 sequences stand out as being clearly more efficient for transactivation (Figure 2C).

To comfort this conclusion, and ascertain that the result can be generalized to other nucleotide contexts, we selected few of the tested DR4, cloned them in a luciferase reporter vector and addressed their capacity to transactivate the reporter gene expression. Nine different DR4 elements were tested confirming the main conclusions of the STARRseq analysis. Interestingly, the response elements differed not only by their capacity to drive luciferase expression upon T3 stimulation, but also by their ability to repress reporter expression. Therefore the DR4 elements ranking differed, depending on the chosen criterion: induction rate or maximal expression in presence of T3 (Figure 3).

**Figure 3:**
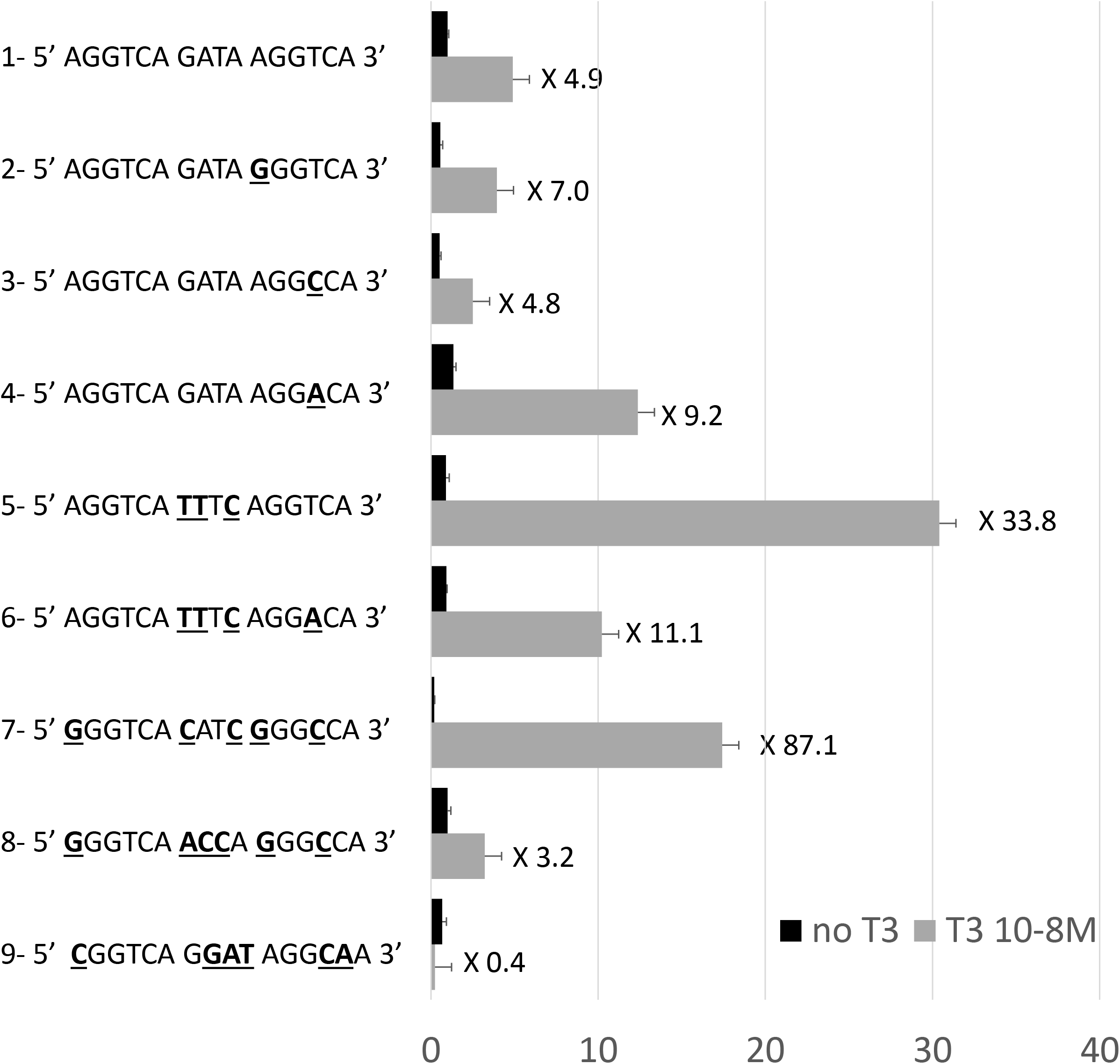
Individual tests of 9 DR4 sequences in transient expression assays. DR4-1 is the most efficient of the tested DR4, according to figure 2C. Comparisons between DR4-1 −2 −3 and −4 highlight the most favorable nucleotide in the 3’ half-site occupied by TRα1. Comparisons with DR4 −5 −6 −7 −8 demonstrate that spacer nucleotides have an important influence on transactivation and that a T at the third position of the spacer (position 9) is more favorable to transactivation. DR4-9 was predicted to be inactive by STARRseq analysis, which is confirmed in this assay. Note that the negative influence exerted by unliganded TRα1, and that induction rate is maximal for DR-7. Two to 4 repeats of DR4-6 were previously used in reporter constructs to provide maximal transactivation capacity (18).

We then addressed the possibility that this functional screen reflects the capacity of specific sequences to bind TRs, as defined by genome-wide analysis of chromatin occupancy. We used for this a recently published Chip-Seq dataset (9) describing chromatin occupancy by TRα1 in the GABAergic neurons of mouse striatum. Using FIMO search, from the MEME suite (17), we listed 51231 matches in the mouse genome (GRCm38/mm10) for the DR4 motif (5’NGGTCANNNNRGGNNA3’) used for our STARR-seq screen. Intervals intersection (Galaxy Version 2.30.0) identified 2099 out of these 51231 motives, which were occupied by TRα1 in this cell type. We then asked whether the nucleotide composition of the occupied and unoccupied DR4 motives differ. Table 2 reveals striking similarities between the outcome of this differential analysis and of the STARR-seq screen. There is therefore a strong correlation between the capacity of a specific DR4 element to recruit TRα1 in chromatin, and the capacity of this DR4 to mediate transactivation in transfected cells. We pursue the comparison between the outcome of ChipSeq and STARR-seq analyses and tested the influence of combination of nucleotides. We also found that some specific combinations at position 14 and 15 are also more favorable to chromatin occupancy by TRα1 (Table 3). With only 2099 TRα1 binding sites in chromatin, this statistical comparison was not applicable to the spacer combinations, or to the whole sequence.

**Table 2:** Relative frequency of nucleotide in DR4 elements occupied by TRα1 according to Chip-Seq. The data are based on a Chip-Seq analysis performed in mouse striatum.

| Position | 1 |  | 7 | 8 | 9 | 10 |  |  | 14 | 15 |  |
| --- | --- | --- | --- | --- | --- | --- | --- | --- | --- | --- | --- |
| Seq 5' to 3' | N | GGTCA | N | N | N | N | A/G | GG | N | N | A |
| A | 1.13 |  | 1.46 | 0.93 | 0.27 | 0.90 | 1.32 |  | 0.94 | 0.42 |  |
| C | 0.37 |  | 0.97 | 1.00 | 1.09 | 1.39 |  |  | 0.65 | 2.11 |  |
| G | 1.97 |  | 0.60 | 1.21 | 0.53 | 1.64 | 0.63 |  | 0.44 | 0.68 |  |
| T | 0.08 |  | 1.74 | 0.87 | 2.54 | 0.41 |  |  | 2.23 | 0.50 |  |

**Table 3:** Associations of nucleotides at position 14 and 15 in DR4 occupied by TRα1. Relative frequency of a given dinucleotide at position 14 and 15 (Occupied by TRα1/Unoccupied). Values above 1.5 are in bold characters. Compare to figure 2A.

|  | Pos.15 | A | C | G | T |
| --- | --- | --- | --- | --- | --- |
| Pos.14 | A | 0.23 | **2.18** | 0.36 | 0.94 |
|  | C | 0.45 | **1.59** | 0.98 | 0.29 |
|  | G | 0.20 | 0.82 | 0.26 | 0.28 |
|  | T | 1.18 | **3.66** | **1.57** | 0.99 |

## Discussion

Our report represents an unprecedented wide analysis of the transactivation capacity of TRα1/RXRα heterodimers. We used an improved version of the synthetic STARR-seq protocol, which proves to reduce background transcription and facilitates the detection of hormonal response. Compared to the previous similar study, performed with the glucocorticoid receptor (15), we observed higher induction rates and enrichments. The protocol described above should thus be directly applicable for other nuclear receptors response elements. In the case of TRα1/RXRα heterodimers, we first identifies 455 efficient DR4 elements, and confirmed in independent transient expression assays that some of them outperform the one previously used in reporter constructs (18).

The main conclusion of our study is the striking concordance between functional tests and chromatin occupancy analyses. Previous low throughput analyses which used naked DNA are also concordant, indicating for example that a 5’TGTA or 5’TGAG spacer is much more favorable than 5’TGAT or 5’TGAC and that the AGGTCA is also optimal for the 5’ half-site (19). We thus conclude that the transactivation capacity of a given DR4 essentially reflects the affinity of the TRα1/RXRα heterodimer for its DNA sequence. As expected from the structural data, the two last-nucleotides of the DR4 spacer participate in defining this affinity.

Only a small subset of the many putative DR4 elements present in the genome are occupied by TRα1/RXRα heterodimers in a given cell type, as judged from Chip-Seq analysis. Despite some variations (unpublished data) a significant part of this cistrome is shared by different cell types, and contains the functional DR4 elements that we identified. By contrast, the repertoire of genes which transcription is activated by T3 is highly dependent on the cell types. We thus did not expect that specific DR4 would be present in the TR binding site associated to the 213 genes which are T3 responsive genes in striatum GABAergic neurons, and this was not the case (data not shown). The explanation to this cell specific response to T3 is thus more likely to be found in the cellular context, for example as a result of variations in the repertoire of transcription coactivators.

## DATA AVAILABILITY

Raw data and processed tables are available at the Gene Expression Omnibus (GSE196145) The following secure token has been created to allow review of record GSE196145 while it remains in private status: iloxqeocttmjxqz

## SUPPLEMENTARY DATA

Supplementary spreadsheet files (Table S1 to S5) are combined in a single MS Excel (.xls)

## ACKNOWLEDGEMENT

We thank Sandrine Hughes and Benjamin Gillet of the PSI facility for technical help and discussions and Sophia Belkhir for gathering preliminary data.

